# Over 2.5 million COI sequences in GenBank and growing

**DOI:** 10.1101/353904

**Authors:** Teresita M. Porter, Mehrdad Hajibabaei

## Abstract

The increasing popularity of cytochrome c oxidase subunit 1 (COI) DNA metabarcoding warrants a careful look at the underlying reference databases used to make high-throughput taxonomic assignments. The objectives of this study are to document trends and assess the future usability of COI records for metabarcode identification. Over 2.5 million COI sequences were found in GenBank, half of which were fully identified to the species rank. From 2003 to 2017, the number of COI Eukaryote records deposited has grown by two orders of magnitude representing a nearly 42-fold increase in unique species. For fully identified records, 92% are at least 500 bp in length, 74% have a country annotation, and 51% have latitude-longitude annotations. To ensure the future usability of COI records in GenBank we suggest: 1) Improving the geographic representation of COI records 2) Improving the cross-referencing of COI records in the Barcode of Life Data System and GenBank to facilitate consolidation and incorporation into existing bioinformatic pipelines, 3) Adherence to the minimum information about a marker gene sequence guidelines, and 4) Integrating metabarcodes from eDNA and mixed community studies with existing sequences. COI metabarcoders are normally considered consumers of taxonomic data. Here we discuss the potential for taxonomists to reverse this pattern and instead mine metabarcode data to guide species discovery. The growth of COI reference records over the past 15 years has been substantial and is likely to be a resource across many fields for years to come.

## Introduction

Cytochrome c oxidase subunit 1 (COI) marker gene or DNA barcode sequencing of animals from mixed communities and bulk samples has surged in usage [1]. It is hoped that COI metabarcoding, a scalable method that can take advantage of automated work-flows, can improve throughput and facilitate large-scale studies [2,3]. COI metabarcoding applications include diversity assessments for biomonitoring and conservation [4,5], detection of environmental gradients in ecology and forestry studies [6,7], and diet analysis [8,9].

COI metabarcoding leverages existing COI sequences in databases such as the Barcode of Life Data (BOLD) System as well as the International Sequence Database Collaboration (INSDC) between the National Center for Biotechnology Information (NCBI), the European Bioinformatics Institute (EMBL-EBI), and the DNA Data Bank of Japan (DDJB) [10,11]. Automated taxonomic assignment of anonymous COI metabarcodes from mixed samples depends on the availability of representative reference sequences for comparison. For taxonomic assignment of large batches of COI sequences from high throughput sequencing platforms, popular methods include the top BLAST hit approach or the naïve Bayesian COI Classifier [12,13]. Both of these methods rely on publically available COI reference sequences. Overviews of the taxonomic coverage of COI sequences in the NCBI nucleotide database have been published [13–15]. Previous work has focused on invertebrates and Insecta for freshwater biomonitoring in general and for Canadian freshwater biomonitoring in particular.

Past studies have looked at the ability of COI metabarcodes to identify taxa from mixed samples by using different taxonomic assignment methods [13,14]. A common assumption is that target taxa are present in the reference database for comparison. More often than is acknowledged, this assumption is false. When target taxa are missing from the reference database, users run the risk of making false positive taxonomic assignments [16]. The mis-assignment of a metabarcode to the wrong species, because the target species is missing from the database has been called a false-positive assignment or over prediction in the literature [17]. The current study takes a step back to outline the type and quality of COI data contained in the NCBI nucleotide database and the implications for future work.

Here we focus on the current level of taxonomic and metadata annotation of COI sequences in the NCBI nucleotide database. We describe trends since the inception of COI barcoding and implications for COI metabarcoding going forward. We highlight two COI metabarcoding applications: 1) freshwater invertebrate biomonitoring, and 2) detection of endangered animal species as listed by International Union for Conservation of Nature (IUCN). We chose these examples to illustrate COI coverage for two very different metabarcoding applications. For freshwater benthic taxa, we anticipated good COI database representation [5,15]. We also hoped to illustrate the potential for COI metabarcoding of endangered animal species based on encouraging results from previous COI barcoding studies of Bovidae, antelopes, and placental mammals [18–20]. Metadata analysis shows that the COI records in the NCBI nucleotide database have increased substantially since the introduction of COI barcoding to the community and includes records with a global geographic distribution. In this high-level analysis, we highlight a few areas to improve COI sequence usability across studies: 1) Improving the representation of COI records from more diverse geographic regions, 2) Improving the cross-referencing of COI records in BOLD and the NCBI nucleotide database to facilitate consolidation and incorporation into existing bioinformatic pipelines, 3) Adherence to the minimum information about a marker gene sequence (MIMARKS) guidelines, and 4) Integrating metabarcodes from eDNA and mixed community studies with fully identified sequences from individual specimens.

## Bioinformatic Methods

GenBank data was parsed using a combination of command-line and custom Perl scripts using BioPerl modules [21]. Tabular data was formatted using Python and plotted in R [22]. We use the terminology from Nilsson et al., (2005) and refer to taxa identified to the species rank as ‘fully identified’ and all other taxa as ‘insufficiently identified’ [23]. We also focused on NCBI nucleotide data deposited from 2003, the year COI barcoding was first introduced to the community, to present (2017) [24].

The names and taxonomic identifications for all Eukaryotes annotated to the species rank were retrieved from the NCBI taxonomy database using the Entrez query “Eukaryota[ORGN]+AND+species[RANK]” with an ebot script [Accessed November 3, 2017] [25]. Taxa were filtered according to the contents of the species field. Only fully identified taxa with a complete Latin binomial (genus and species) were retained. Entries that contained the abbreviations sp., nr., aff., or cf. were discarded. The remaining species names were formatted for use in the next query [*species list*]. For each year from 2003 – 2017 [*year*], records in the NCBI nucleotide database containing COI sequences were retrieved using the Entrez query “(“CO1”[GENE] OR “COI”[GENE] OR “COX1”[GENE] OR “COXI”[GENE]) AND “Eukaryota”[ORGN] AND [*year*][PDAT]) AND [*species list*]” [2003 – 2016, accessed November 2017; 2017, accessed April 2018]. GenBank records were parsed, retaining information on year of record deposition and number of fully identified records. For fully identified records, sequence length as well as country and/or latitude-longitude fields were parsed.

We also assessed the number of high quality COI sequences that meet the standards developed between the INSDC and the Consortium for the Barcode of Life by looking for the BARCODE keyword in the GenBank record [11]. For each year from 2003 – 2017 [*year*], records in the NCBI nucleotide database containing COI BARCODE sequences were retrieved using the Entrez query “(“CO1”[GENE] OR “COI ”[GENE] OR “COX1”[GENE] OR “COXI”[GENE]) AND “Eukaryota”[ORGN] AND [*year*][PDAT] AND “BARCODE”[KYWD]) AND [*species list*]”. Fully identified and geotagged records were parsed as described above.

For our application example on freshwater biomonitoring indicator taxa, we retrieved a high-level list of target taxa from Elbrecht and Leese (2017) [26]. Target freshwater taxa included: Annelida classes Clitellata and Polychaeta; Insecta (Arthropoda) orders Coleoptera, Diptera, Ephemeroptera, Megaloptera, Odonata, Plecoptera, and Trichoptera; Malacostraca (Arthropoda) orders Amphipoda and Isopoda; Mollusca classes Bivalvia and Gastropoda; and Platyhelminthes class Turbellaria. For each freshwater target group we queried the NCBI taxonomy database for records identified to the species rank as described above. These taxon ids were concatenated and used to query the NCBI nucleotide database as described above. We assessed the representation of freshwater indicator taxa in the NCBI nucleotide database and level of annotation as described above.

For our application example on IUCN endangered animal species, we retrieved a list of endangered species names from http://www.iucnredlist.org from all available years (1996, 2000, 2002-2004, 2006-2017) filtering the results for native Animalia species [Accessed Dec. 12, 2017]. We excluded insufficiently identified species containing the terms ‘affinis’, ‘sp.’, or ‘sp. nov.’, leaving us with a list of 4,289 endangered animal species as well as 2,089 synonyms. We submitted this combined list of species names to the ‘NCBI Taxonomy name/id Status Report Page’ (https://www.ncbi.nlm.nih.gov/Taxonomy/TaxIdentifier/tax_identifier.cgi) and retrieved a list of 2,613 taxon ids. For each taxon id, we queried the NCBI taxonomy and nucleotide databases as described above.

To assess the number of COI records unique to the BOLD database compared with the NCBI nucleotide database, we also retrieved records from the BOLD Application Programming Interface (API) as well as from the data releases. COI sequences were retrieved from the BOLD API (http://www.boldsystems.org/index.php/API_Public/sequence?) using the terms ‘marker=COI-3P|COI-5P&taxon=’ for each Eukaryote phylum except for Arthropoda which was queried separately for each class, and Insecta which was queried separately for each order to enable the download of complete files [Accessed Apr. 26, 2018]. Lists of Eukaryote phyla, Arthropoda classes, and Insecta orders were retrieved from the BOLD taxonomy browser (http://www.boldsystems.org/index.php/TaxBrowser_Home). COI records were also retrieved from the BOLD data releases (http://www.boldsystems.org/index.php/datarelease). All available releases of animal COI records up to and including Release 6.50v1 were individually downloaded and parsed. Note that the records retrieved from the data releases may not be as current as those retrieved through the BOLD API.

## Results

The dataflow and scripts used in this study are available from GitHub (xxx).

### COI record growth in GenBank

A total of 2,530,418 COI Eukaryote records were identified from the NCBI nucleotide database. Of these, 1,383,206 (55%) records were verified to be fully identified COI records. We show the growth of COI records in GenBank from the introduction of COI barcoding in 2003 to present (2017). The number of COI records deposited to GenBank increased from 8,137 in 2003 to 2,522,281 from 2004-2017, an increase of two orders of magnitude (Fig 1). This corresponds to a nearly 42-fold increase in the number of unique species with COI sequences deposited during this time. Since 2010, the proportion of insufficiently identified COI records has also grown substantially compared with the number of fully identified records.

**Fig 1:**
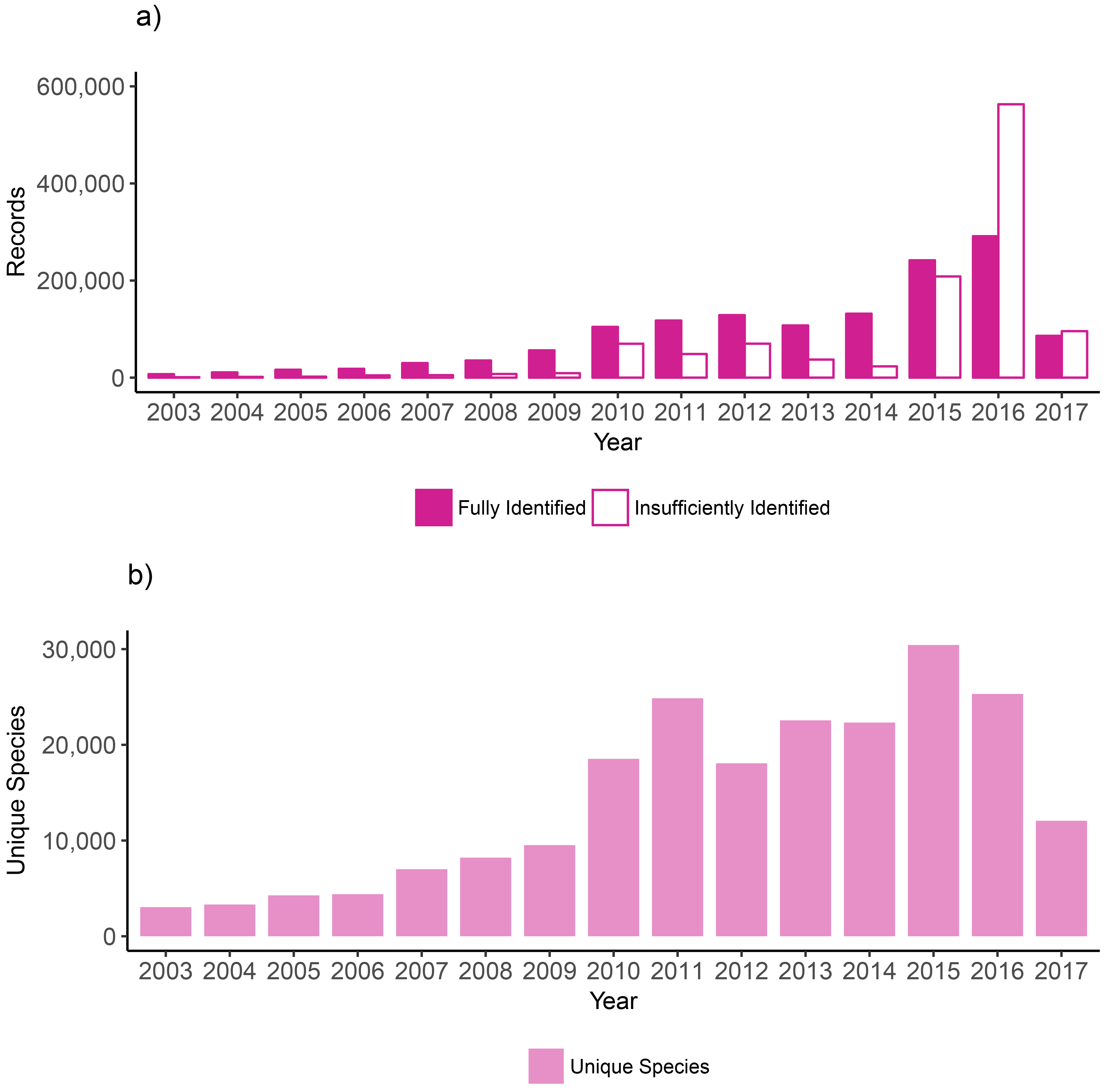
The number of Eukaryote COI records and unique species in the NCBI nucleotide database has grown since 2003. A) The number of records deposited from 2003 to 2017. B) The number of unique species that the fully identified records represent.

### Target COI taxa in the NCBI nucleotide database

Large numbers of COI Eukaryote NCBI nucleotide records can be described as BARCODE or freshwater biomonitoring taxa (Fig 2). 718,814 (28%) are flagged with the BARCODE keyword indicating these records meet the standards created in consultation with Consortium for the Barcode of Life [11]. 1,096,518 (43%) represent high-level freshwater biomonitoring taxa of interest. Records for freshwater taxa largely represent Diptera (true flies, 728,906), Coleoptera (beetles, 151,841), and Gastropoda (snails and slugs, 76,786) (S1 Figure). 1,190 (28%) of the IUCN endangered animal species have corresponding COI records in GenBank. A total of 11,934 NCBI nucleotide COI records correspond to IUCN endangered animal species (S2 Figure).

**Fig 2:**
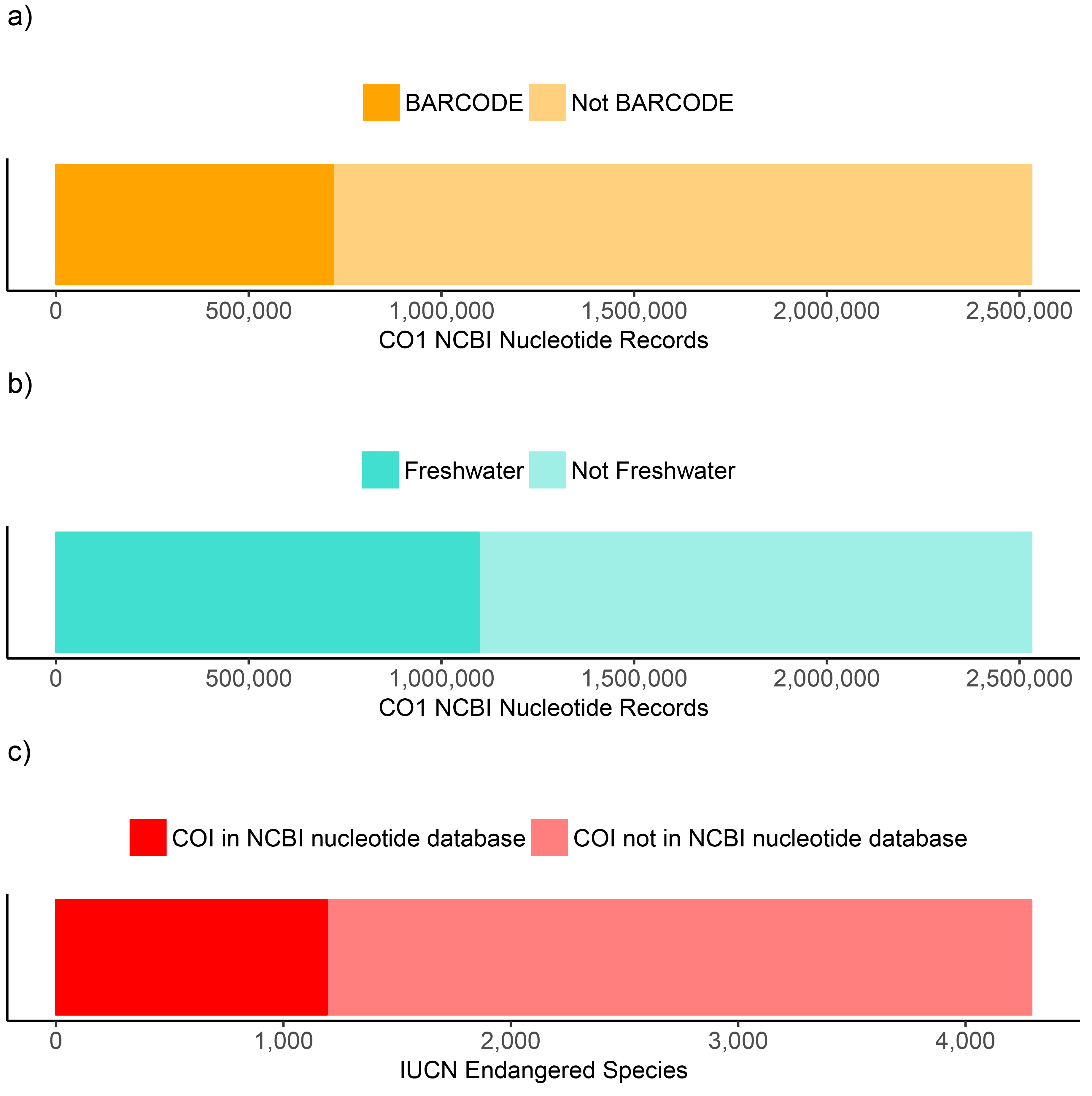
Summary of target taxonomic groups in the NCBI nucleotide database. A) The proportion of all Eukaryote COI records flagged with the BARCODE keyword. B) The proportion of all Eukaryote COI records that represent high-level freshwater biomonitoring target taxa. C) The proportion of IUCN endangered animal species that are represented by COI records.

We also show that the number of COI records deposited to the NCBI nucleotide database for specific groups of taxa (BARCODE, freshwater, endangered) has substantially grown from 2003 to 2017 (S2 Figure). The number of BARCODE taxa deposited was 386 in 2004 and 718,328 from 2005-2017. The number of freshwater taxa deposited was 3,219 in 2003 and 1,093,301 from 2004-2017. The number of deposited BARCODE and freshwater COI records increased by three orders of magnitude. The number of records deposited that represent endangered species was 402 in 2003 and 11,532 in 2004-2017 representing an increase of two orders of magnitude or nearly 29-fold.

### COI NCBI nucleotide record annotations

Overall COI record annotation completeness was highest for BARCODE flagged records (Fig 3). The proportion of fully identified BARCODE records was 51% and similar to the level of fully identified records for All Eukaryotes and the subset of freshwater taxa. Nearly all of the fully identified BARCODE records had good sequence length (500 bp+) and were geotagged with country and latitude-longitude information. In contrast, the proportion of endangered species that were fully identified is 100% by default because we were searching for a specific list of endangered species. Records for endangered species were relatively incomplete with 49% that included country and 18% that included latitude-longitude data.

**Fig 3:**
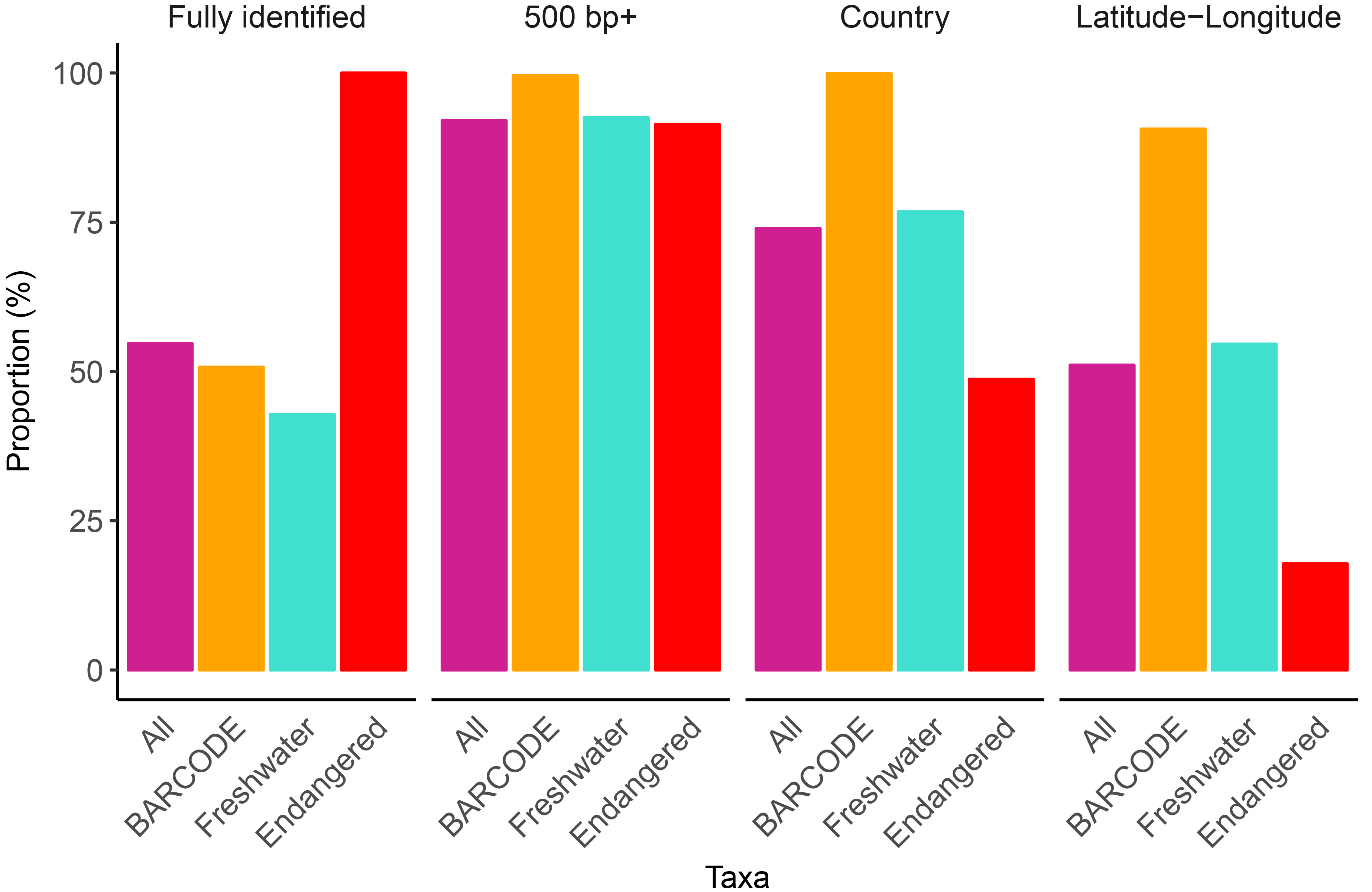
COI BARCODE records in the NCBI nucleotide database are well annotated. The first panel shows the proportion of records that are fully identified to the species rank. The remaining three panels show the proportion of fully identified records with good sequence length (500 bp +) and are geotagged (country and/or latitude-longitude). ‘All’ refers to the complete set of COI Eukaryote records deposited to the NCBI nucleotide database from 2003 to 2017. BARCODE refers to the subset of records flagged with the BARCODE keyword. Freshwater refers to the subset of records that represent high-level freshwater biomonitoring taxa of interest. Endangered refers to the subset of fully identified records on the IUCN endangered animal species list.

### Geographic distribution of COI records

Fully identified COI NCBI nucleotide records show a global distribution but are biased towards Canada (364,356) (Fig 4). There are nearly as many fully identified records with no country data provided (364,356). The 5 next best-represented countries are USA (78,121), Costa Rica (46,597), Australia (41,019), China (36,250), and Germany (34,864). Although country annotation data are useful, because of variations in spelling, as well as country borders and names that change over time, this can be a difficult metadata field to standardize across studies. Latitude-longitude data provide more resolution of CO1 record distribution within countries and are easier to combine across data sets but we found this data is often lacking in non-BARCODE COI records. Similar maps for BARCODE, freshwater, and endangered animal species are also provided (S3 Figure).

**Fig 4:**
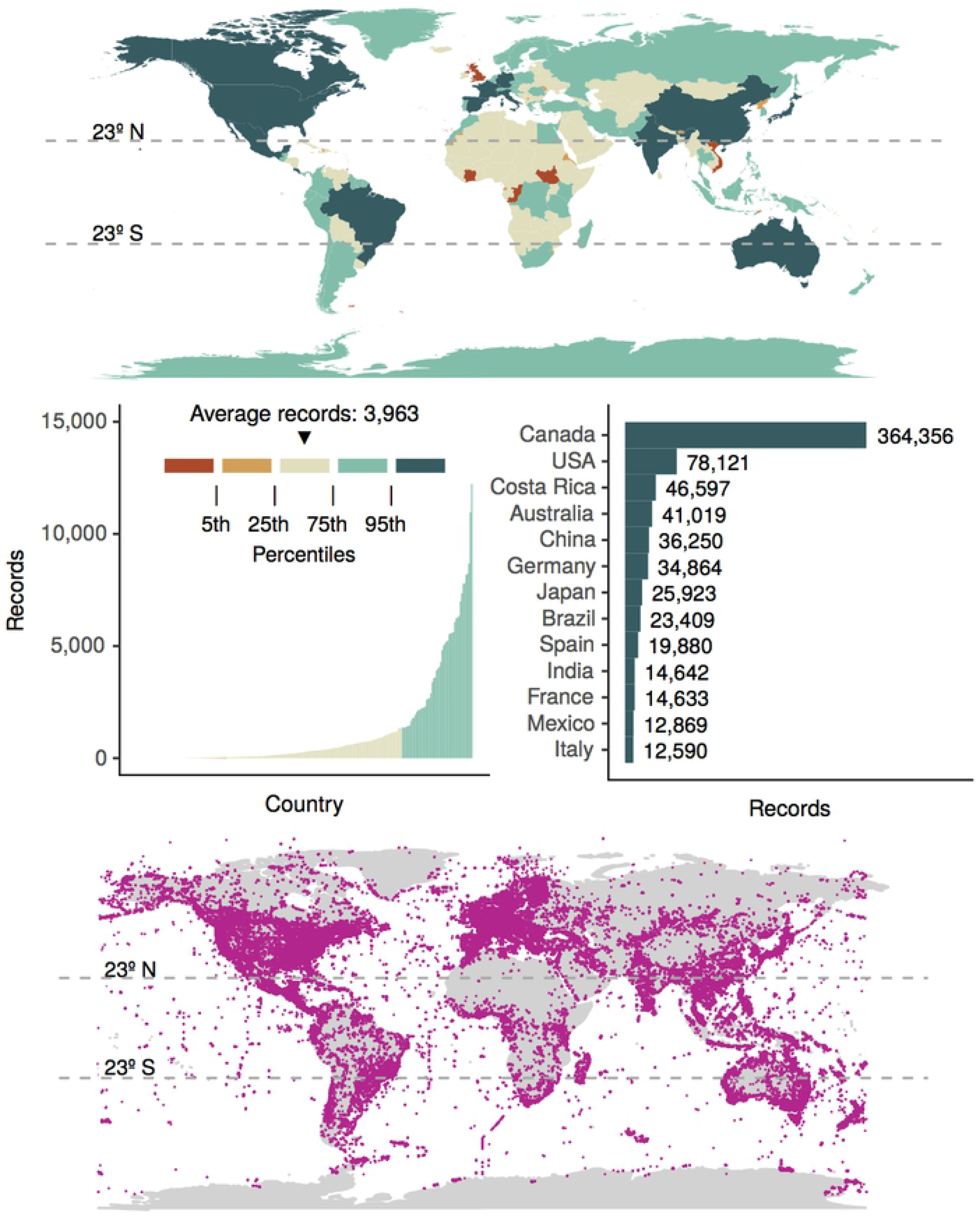
Worldwide distribution of fully identified COI Eukaryote NCBI nucleotide records. Top panel: The number of records per country, where this information was available, is shown. Middle panel: The number of records per country with the bottom 95% of countries shown on the left, and the top 5% of countries shown on the right. Bottom panel: Latitude-longitude data, where this information was available, are plotted as pink points.

After filtering out data from non-COI markers, we retained a final set of 4,646,172 COI sequences from the BOLD API (Table 1). 15% (705,711) of these sequences were associated with a GenBank record flagged with the BARCODE keyword. 48% (2,238,104) had GenBank accessions that were not flagged with the BARCODE keyword. This set of the records seems to be either mined from GenBank and used to supplement the BOLD database, or they were BOLD records deposited to GenBank then subsequently suppressed because the record did satisfy the iBOL/GenBank early release agreement. The remaining 37% (1,715,362) of the records appear to be unique to BOLD.

**Table 1:** Distribution of COI records in BOLD

| **BOLD Source** | **BOLD Record with GenBank accession + BARCODE keyword** | **BOLD Record with GenBank accession** | **Remaining BOLD Records** | **Total** |
| --- | --- | --- | --- | --- |
| BOLD API | 705,711 (15%) | 2,238,104 (48%) | 1,715,362 (37%) | 4,646,172 |
| BOLD Data Releases | 341,150 (13%) | 896,150 (33%) | 1,490,534 (55%) | 2,727,834 |

A total of 2,727,834 unique records were retrieved from the BOLD data releases (Table 1). 13% (341,150) were associated with a GenBank record flagged with the BARCODE keyword. 33% (896,150) were associated with a GenBank record but did not contain a BARCODE keyword. Some of these records were found to contain iBOL details in the GenBank record COMMENT section, others were not indicated as being BOLD records in any way. 55% (1,490,534) of the records appear to be unique to BOLD. Whether or not a linked GenBank record was subsequently suppressed is not indicated here.

## Discussion

### Taxonomic representation

Based on existing work, we already knew that the best-represented taxonomic groups of COI sequences in the NCBI nucleotide database were Arthropods, followed by Chordata [13]. More specifically, Insecta and Diptera have the largest number of COI records in GenBank [14]. But is this data suitable for real-world application of COI metabarcoding for freshwater biomonitoring? A Canadian study by Curry et al. (2018) looked specifically at taxa expected to be recovered during freshwater biomonitoring using traditional methods and checked for their presence in GenBank and BOLD [15]. They found that ~ 61% of North American freshwater genera were associated with COI sequences in both databases, but either database alone provided coverage of ~51% of genera. Practically, however, they found that ~95% of genera found in more than 1% of field samples were associated with COI sequences. In the current study, we found that ~ 43% of the COI records in the NCBI nucleotide database represent high-level freshwater taxa of interest for biomonitoring. Similar to Curry et al., 2018, we also found here that Diptera are especially well-represented in GenBank [15]. When we look at the quality of the COI freshwater records in the NCBI nucleotide database, we found that ~ 43% were fully identified to the species rank. If trends continue, the number of unique species represented by COI records will continue to improve with time. With this in mind, reanalyzing older data as public databases are updated should improve the proportion of metabarcodes that are identified.

### Geographic representation

Having a reference database that represents local species is important for making reliable COI metabarcode assignments. As shown by the Curry et al., (2018) as well as the current study, a good proportion of GenBank COI records represent Canadian freshwater biomonitoring taxa of interest. For studies outside of Canada, or for applications other than freshwater biomonitoring, an assessment of whether target taxa are represented in current databases (BOLD or GenBank) should be performed prior to conducting taxonomic assignments. Understanding current database composition, where the gaps exist, can help guide future work, i.e., interpreting the results from bioinformatics pipelines, choosing the level of reported taxonomic resolution, and determining statistical confidence for taxonomic assignments [13]. Specific taxonomic gaps in COI GenBank data have been previously published and is beyond the scope of this study [13–15]. The current study highlights both the global distribution of COI records in GenBank, as well as the variability in geographic representation for different subsets of the data, i.e. freshwater taxa of interest versus endangered animal species. We show here that some areas of the world known to have very high endemic diversity, i.e. the tropics [27,28], are disproportionally under-represented by fully identified COI records.

### BOLD data in the NCBI nucleotide database

COI records in BOLD and the NCBI nucleotide database are not fully synced or consistently cross-referenced. We show here that a significant portion of the COI records in GenBank, 718,714 (~ 28%) have been flagged with the BARCODE keyword. These records represent 13-15% of COI sequences retrieved from BOLD. We found that a further 33-48% of BOLD records are associated with GenBank records but that these GenBank records are inconsistently cross-referenced with BOLD records (lacking the BARCODE keyword, or cross referencing information placed in the comments field) or they have been subsequently suppressed in GenBank for technical reasons. If the community could improve the cross-referencing of BOLD and GenBank records this could facilitate the re-usability of COI records across studies.

37 - 55% of BOLD records may be unique to BOLD. It is this subset of the BOLD data that users who create custom databases don’t want to miss. Users are often faced with the choice of using either BOLD or INSDC data for identification by creating custom COI databases to permit high-throughput identifications. The recently published BOLD_NCBI_MERGER script helps to combine records from BOLD with those in GenBank for use with BLAST and MEGAN lowest common ancestor taxonomic assignment [29]. The tool helps to combine high quality COI barcode records from BOLD with the broader taxonomic coverage of COI records from the NCBI nucleotide database. Future taxonomic assignment method developments would likely benefit from combining these databases to improve overall COI record representation.

### Minimum information about marker gene sequences (MIMARKS)

The Genomics Standards Consortium (GSC) has already outlined recommendations for the minimum information about a marker gene sequence (MIMARKS) that should be submitted with released sequences [30]. That study indicates which metadata fields should be mandatory or environment-specific. Whenever possible, values are based on a controlled vocabulary or ontology. Major databases such as BOLD and GenBank already support these standards. We show here that across the COI Eukaryote NCBI nucleotide records 74% have country and 51% have latitude-longitude metadata (part of MIMARKS). In contrast, nearly all GenBank BARCODE records have country and latitude-longitude metadata. If the community could further improve their compliance with MIMARKS this could greatly contribute to the re-usability of COI GenBank data across studies.

### Incorporating COI references into existing bioinformatics pipelines

The significance of metabarcoding for ecology and biomonitoring more broadly have been shown [31]. COI metabarcoding has its roots in the COI barcoding initiative as well as the long history of metagenomic marker gene surveys for microbial ecology investigations [32–34]. Reference databases such as SILVA and GreenGenes for 16S rDNA as well as UNITE for ITS rDNA [35–37] have been integrated into popular bioinformatics pipelines such as MOTHUR and QIIME2 (https://qiime2.org/) [38,39]. To enable COI sequences to be analyzed along-side other popular metabarcoding markers, future work should make curated COI reference sets available to the broader community through similar popular bioinformatics pipelines that have already been widely adopted.

### The problem of insufficiently identified sequences: hidden opportunities

It is not uncommon for current COI metabarcode studies to only identify a fraction of the total number of sequences, operational taxonomic units, or exact sequence variants. It has been assumed that the remaining sequences represent a mix of sequence artefacts (non-specific amplification products, chimeric sequences, sequencing errors, etc.) and real species that remain insufficiently identified due to a lack of representatives in reference databases. We have shown in this study that the intersection of COI sequences in BOLD and GenBank is relatively small and this could be one possible reason for insufficiently identified records. On the other hand, if insufficiently identified sequences represent existing named species not present in any public database, then sequencing type specimens should improve taxonomic assignment rates. Similar initiatives have already been initiated for prokaryotes and fungi [40–42]. If, however, insufficiently identified sequences from metabarcode studies represent new taxa, this implies that metabarcoding studies may also be an important new tool for local species discovery as has been found for prokaryotes and fungi [43,44]. This distinction is important because until now COI metabarcoders have been consumers of taxonomic information, such as the high quality records provided by BOLD. It may now be possible for taxonomists to turn the table on this relationship and mine metabarcode data for novel species. In conjunction with non-destructive sampling methods, vouchered bulk samples (e.g. from benthic kicknets) could harbor intact new specimens for formal description using more traditional methods [45,46]. Geotagged records could also guide taxonomists on where to search for novel local taxa.

Another way to handle insufficiently identified sequences, for example from eDNA and mixed community studies, would be to integrate them with existing fully identified sequences. With fungal ITS rDNA, for example, an increasing proportion of insufficiently identified sequences was documented along-side the rise in use of DNA-based methods for ecological studies [47,48]. The explosion in insufficiently identified fungal ITS rDNA sequences effectively out-paced the ability for traditional taxonomy to study and name all new species. Instead, the disambiguation of insufficiently identified sequences was addressed by developing species hypotheses (SH) [49]. The SH concept is similar to a COI Barcode Index Number (BIN) in that it is a cluster of similar sequences, and each SH is given a stable numeric label in the UNITE database. Fungal SH’s takes the BIN concept a step further by clustering insufficiently identified sequences from *environmental samples* into clusters with stable naming to allow cross-referencing across other metabarcode studies.

A known issue with clusters, however, is that the composition can change depending on order of sequences in a file (when using greedy clustering methods) or by the clustering algorithm chosen (single-, complete-, average-linkage). A method developed to overcome these issues to generate stable operational taxonomic units (OTUs) is SWARM [50]. In any case, users are left with the difficulty of interpreting their clusters as they may not represent unique species. In this study, we show that the rate of insufficiently identified COI records deposited to GenBank is increasing faster than the rate of fully identified records. Looking forward, we must as a community find a realistic way to integrate fully identified as well as insufficiently identified COI records from all sources including COI barcodes as well as sequences from eDNA and mixed community studies.

An emerging practice in the metabarcoding/marker gene community has been to move away from working with sequence clusters and instead focus on exact sequence variants (ESVs) [51]. An ESV can be thought of as an OTU defined by a sequence similarity cutoff of 100%. ESVs have a more straight-forward interpretation than OTUs, which can facilitate easier combinability across studies. New COI sequences can simply be mapped to existing ESVs with 100% sequence similarity and remaining unique sequences become new ESVs. Another benefit is improved resolution by avoiding the accidental ‘lumping’ of ESVs from different species into single clusters [52]. We can envisage how COI ESVs, generated from individual specimens as well as eDNA and mixed community studies, could be combined with a stable numbering system to allow for standardized cross-referencing. Such a method would allow for the detection of biodiversity at a finer level of resolution, capturing sequence-level variation and geographic patterns that would otherwise be obscured in BIN clusters.

We have demonstrated the growth of COI reference records over the past 15 years. We have emphasized the importance of including geographic metadata with COI sequences deposited to the INSDC. Growth in the adoption of COI metabarcoding applications has been substantial in recent years making high quality public COI reference databases an important resource across many fields for years to come.

## Acknowledgements

The authors would like to thank the Canadian government for funding through the Genomics Research and Development Initiative (GRDI) interdepartmental EcoBiomics project.

## Supporting Information Captions

S1 Figure. The number of CO1 GenBank records for freshwater biomonitoring target taxa in the NCBI nucleotide database are biased towards Diptera.

S2 Figure. The number of CO1 GenBank records deposited in the nucleotide database has grown since 2003. A) The CO1 barcoding initiative was first introduced by Hebert et al. (2003) and the first CO1 records flagged with the BARCODE keyword were deposited in 2004 [24]. B) The number of records deposited for freshwater biomonitoring target taxa were tracked from 2003 to 2017. C) The number of records that represent IUCN endangered species were tracked from 2003 to 2017.

S3 Figure. Fully identified CO1 GenBank records have a global distribution that varies according to data partition. The number of records per country, where this data is available, is shown in the legend (log scale): A) BARCODE, B) Freshwater, C) IUCN endangered species. Latitude-longitude data, where this data is available, is plotted as points in ‘orange’ for BARCODE records, in ‘turquoise’ for freshwater records, and in ‘red’ for endangered animal species.

